# Structure-function analysis of the maize bulliform cell cuticle and its role in dehydration and leaf rolling

**DOI:** 10.1101/2020.02.06.937011

**Authors:** Susanne Matschi, Miguel F. Vasquez, Richard Bourgault, Paul Steinbach, James Chamness, Nicholas Kaczmar, Michael A. Gore, Isabel Molina, Laurie G. Smith

## Abstract

The cuticle is a hydrophobic layer on the outer surface plant shoots, which serves as an important interaction interface with the environment. It consists of the lipid polymer cutin, embedded with and covered by waxes, and provides protection against stresses including desiccation, UV radiation, and pathogen attack. Bulliform cells form in longitudinal strips on the adaxial leaf surface, and have been implicated in the leaf rolling response observed in drought stressed grass leaves. In this study, we show that bulliform cells of the adult maize leaf epidermis have a specialized cuticle, and we investigate its function along with that of bulliform cells themselves. Analysis of natural variation was used to relate bulliform strip pattering to leaf rolling rate, providing evidence of a role for bulliform cells in leaf rolling. Bulliform cells displayed increased shrinkage compared to other epidermal cell types during dehydration of the leaf, providing a potential mechanism to facilitate leaf rolling. Comparisons of cuticular conductance between adaxial and abaxial leaf surfaces, and between bulliform-enriched mutants vs. wild type siblings, provided evidence that bulliform cells lose water across the cuticle more rapidly than other epidermal cell types. Bulliform cell cuticles have a distinct ultrastructure, and differences in cutin monomer content and composition, compared to other leaf epidermal cells. We hypothesize that this cell type-specific cuticle is more water permeable than the epidermal pavement cell cuticle, facilitating the function of bulliform cells in stress-induced leaf rolling observed in grasses.

**One sentence summary:** Bulliform cells in maize have a specialized cuticle, lose more water than other epidermal cell types as the leaf dehydrates, and facilitate leaf rolling upon dehydration.

## Introduction

Plants display a variety of responses to environmental stresses. An important drought stress response in grasses is reversible leaf rolling along the longitudinal leaf axis upon water limitation or heat stress conditions. Leaf rolling prevents water loss, photosynthetic loss, and increases drought resistance in numerous species of the Poaceae (Kadioglu and Terzi, 2007; Saglam et al., 2014), which include staple crops like wheat, rice and maize. Dehydrated grass leaf blades fold longitudinally, reducing the exposed leaf surface area, with effect on leaf transpiration and canopy temperature (O’Toole et al., 1979; Turner et al., 1986). Leaf rolling is linked to osmotic adjustment and a change in leaf water potential upon dehydration (O’Toole and Cruz, 1980; Hsiao et al., 1984; Moulia, 1994). Changes in concentrations of organic acids or ions, accumulation of phytohormones, in part followed by changes in stress-responsive gene expression, and other abiotic and biotic factors contribute to leaf-rolling (Kadioglu et al., 2012).

Bulliform cells (BCs) are enlarged, colorless cells located in the epidermis, which in maize are usually arranged in 2 to 5 cell-wide strips along the longitudinal leaf axis solely on the adaxial side of the leaf (Ellis, 1976; Becraft et al., 2002; Sylvester and Smith, 2009). Being absent in juvenile maize leaves, their emergence in the leaf epidermis is one of the hallmarks of the vegetative phase change in maize, the transition from juvenile to adult leaves (Poethig R. S., 1990). The function of BCs, also called hinge or motor cells, is a matter of ongoing debate since the first description by Duval-Jouve (1875) postulating reversible rolling of the leaf blade resulting from changes in the turgor pressures of BCs. Adaxial rolling is conferred by differential top–bottom elastic shrinkage in the leaf cross-section (Moulia, 2000) which is thought to be guided by BCs due to their asymmetric location on the adaxial leaf surface. However, studies on leaf rolling in different grass types reach contradictory conclusions and could not resolve whether BCs cause rolling by collapse due to water loss, or whether their size and plasticity merely permit them to be compressed and allow rolling to occur (reviewed in Ellis, 1976; Moulia, 2000; Evert, 2006). Numerous mutants with altered BC number, size or adaxial/abaxial patterning, mostly identified in rice and to a lesser extent in maize, show an effect on leaf rolling and underline the importance of this cell type and its distribution for the leaf rolling response (Nelson et al., 2002; Xu et al., 2018; Gao et al., 2019). Only recently the genetic diversity of maize in combination with association mapping and machine learning techniques was employed to characterize the genetic basis of bulliform strip architecture, namely strip number and width, in a collection of diverse maize inbred lines, and identify a number of underlying candidate genes (Qiao et al., 2019).

The cuticle is the first layer of protection of aerial plant tissue and represents an important interaction surface with the environment. It shields the underlying tissue against environmental stresses such as non-stomatal water loss (Riederer and Schreiber, 1995), UV radiation (Krauss et al., 1997) and pathogen attack (Serrano et al., 2014), which in turn can affect cuticle biogenesis (Yeats and Rose, 2013). The biological functions of cuticles are determined by their physical properties, which depend on their chemical composition. The cuticle is comprised of two major components: the polyester cutin, which serves as the cuticle’s framework (Fich et al., 2016), and cuticular waxes, a multitude of mainly hydrophobic compounds that are either embedded into the cutin polymer matrix (intracuticular waxes) or cover the cutin layer (epicuticular waxes) (Koch and Ensikat, 2008). Cutin polymer structures in different species and tissues remain elusive so far, as biochemical characterization of cutin can only be achieved by a description of its monomers after depolymerization. Typical cutin monomers are long-chain (C_16_ and C_18_) fatty acid monomers that have a hydroxy group at the ϖ-position and midchain hydroxy or epoxy groups. Additionally, unsubstituted fatty acids, dicarboxylic acids (DCA), glycerol, and low amounts of phenolic compounds (e.g. hydroxycinnamic acids (HCAs) like coumarate and ferulate) can be present (Pollard et al., 2008). Cuticular waxes are derived from very long-chain fatty acids (VLCFAs), and usually consist of aldehydes, primary and secondary alcohols, hydrocarbons, ketones, and wax esters, and cyclic compounds including terpenoids and sterols (Yeats and Rose, 2013). Unlike cutin, waxes can be extracted from the cuticle with organic solvents.

The cuticle is generally described as having three layers of overlapping composition: 1) the innermost cuticular layer, continuous with the cell wall, which consists of polysaccharides, waxes and cutin; 2) the cuticle proper as the middle layer, with intracuticular waxes embedded in the cutin matrix but devoid of polysaccharides; 3) the epicuticular wax layer as the outermost layer of the cuticle on the plant surface which may be deposited as an amorphous film or in the form of epicuticular wax crystals (Bargel et al., 2006; Jeffree, 2006). Recent reassessments of the literature and emerging techniques however challenge the concept of the cuticle as a lipid layer independent from the cell wall with very defined structural elements (Fernández et al., 2016), and rather describe the cuticle as a form of lipidic modification of the cell wall (Yeats and Rose, 2013). The thickness, structure and chemical composition of cuticular matrices and epi- and intracuticular waxes vary widely between different organisms, developmental stages, and even organs within a species (Jeffree, 2006; Jetter et al., 2006). Understanding how the features of cuticle organization are related to its composition and function is an active area of research.

Retaining water in the epidermis and the underlying plant tissue to limit dehydration is one of the most important roles of the plant cuticle. It is well known that waxes, rather than cutin, provide the majority of the water retention capacity of the cuticle (Schönherr, 1976; Kerstiens, 1996; Isaacson et al., 2009; Jetter and Riederer, 2016). Interestingly, cuticle thickness is no indicator of its function as a water barrier, but wax composition appears to be critical (Buschhaus and Jetter, 2012; Jetter and Riederer, 2016). In a direct comparison of cuticle permeability in several Arabidopsis mutants with either increased or decreased wax and/or cutin loads, most mutants displayed higher permeability than wild-type, even if their respective wax or cutin load was increased or showed a thicker cuticle (Sadler et al., 2016). Increased wax load in cutin mutants is often interpreted as a compensatory mechanism to ensure cuticular integrity despite the insufficient cutin scaffold provided in these mutant backgrounds (Kurdyukov et al., 2006; Bessire et al., 2007).

The overwhelming majority of cuticle studies in maize focus on juvenile leaves, which are, in addition to other distinctive features, quite different in cuticle structure and composition compared to adult leaves (Bianchi and Marchesi, 1960; Bianchi and Avato, 1984; Bongard-Pierce et al., 1996). Only limited research is available on the cuticle composition of mature adult maize leaves and their functional impact, despite the fact that drought stress is most damaging to maize grain yield at the flowering stage (Grant et al., 1989), a time when juvenile leaves have died and only adult leaves remain. The wax profile of the adult leaf cuticle reveals high proportions of wax esters and alkanes and low abundance of free alcohols and aldehydes (Bianchi and Avato, 1984; Bourgault et al., 2020). The cutin polyester in the adult maize leaf mainly consists of di-hydroxy-hexadecanoic acid and typical members of the C_18_ family of cutin acids, including hydroxy and hydroxy-epoxy acids, with low amounts of the HCA derivatives coumarate and ferulate also present (Espelie and Kolattukudy, 1979; Bourgault et al., 2020). In a developmental analysis of the maize leaf cuticle, the establishment of the water barrier properties of the adult leaf cuticle in maize coincided with a switch from alkanes to esters as the major wax type and the emergence of an osmiophilic (likely cutin-rich) layer of the cuticle proper (Bourgault et al., 2020). Ultrastructurally, pavement cell cuticles of the adult leaf did not show a typical three-layered composition, consisting only of a cuticle proper and an epicuticular layer (Bourgault et al., 2020).

Little is known about variation in cuticle composition and structure on specific epidermal cell types and how this relates to their specialized functions. This is challenging to investigate due to the difficulty of separating different epidermal cell types for biochemical analysis of cuticle composition. One example of a comparative study was conducted on trichome cuticles in Arabidopsis (Hegebarth et al., 2016), where comparison of trichome-enriched and depleted genotypes revealed an increase in longer-chain alkanes and alkenes in trichome-rich material. Integration of gene expression data lead to the conclusion that trichomes possess autonomous wax biosynthesis (Hegebarth et al., 2016; Hegebarth and Jetter, 2017). The exposed position of trichomes requires increased flexibility to withstand mechanical stress, but the role of the trichome cuticle, and its specific composition, is unclear (Hegebarth and Jetter, 2017). Furthermore, there is indirect evidence that the cuticles of guard cells have a different composition compared to those of pavement cells but these differences have yet to be characterized (Hegebarth and Jetter, 2017).

The current study aimed to develop a structure-function relationship of the unique BC cuticle with the proposed role of these cells in the leaf rolling response in grasses. Maize’s genetic diversity was used to investigate the impact of architectural features of bulliform strip distribution on leaf rolling speed. On a cellular level, postulated increased water loss of BCs was examined *in situ* with a new cryo-confocal method and dehydration analysis of different BC-enriched tissue types. Variations in cuticles of different epidermal cell types were studied by TEM and lipid staining, and the biochemical composition of several bulliform-enriched or - depleted cuticle types was analyzed to identify a relationship between chemical monomer composition, cuticle ultrastructure and the proposed function of BCs in leaf rolling. Transcriptional analysis of the maize leaf cuticle maturation zone of bulliform-enriched mutants was carried out to identify key players of bulliform cuticle regulation. Together, these experiments provide new insights into the scarce knowledge about cell type-specific cuticles and how structural and compositional cuticle features could connect to the biological function of a specific cell type.

## Results

### Maize leaf rolling is impacted by variation in bulliform strip patterning

Bulliform cells (BCs) are organized into 2-5 cell wide strips aligned with the proximodistal axis of the maize leaf, which are implicated to play a role in the leaf rolling response due to their asymmetric location on the adaxial side of the leaf. To investigate the role of specific architectural features like bulliform strip distribution, e.g. thickness and number, in the leaf rolling response in maize, phenotypic diversity in a panel of diverse maize inbred lines was used to query the relationship between bulliform cell pattern and the rolling response. Adult leaves of maize inbred lines differ dramatically in the frequency and width of bulliform strips (examples in Figure 1A-D, (Qiao et al., 2019)). In addition to their bulliform cell patterns, leaf rolling responses during dehydration were recorded for each of several hundred different maize inbred lines. Adult leaves were excised from each plant, hung from a line to allow them to dehydrate, and scored for their leaf rolling status as a function of time (Supplemental Table S1). Lines with extreme rolling behavior - “fast rollers” (scored as rolled within the first 90 minutes of dehydration, 25 lines), and “never rollers” (no rolling observed in the assessed time frame of 225 minutes, 41 lines) - were selected for analysis. Rolling behavior in these lines was significantly related to the pattern of bulliform cells (Figure 1E-G). Fast rolling leaves had a higher number of BC strips per unit area than never rolling leaves (Figure 1E), while the width of individual bulliform strips showed the inverse trend, with never rolling leaves exhibiting wider strips than fast rolling leaves (Figure 1F). However, overall bulliform coverage was not different between fast and never rollers (Figure 1G). Bulliform strip number and width appear to impact leaf rolling independently, since no correlation was seen between these features across the entire set of inbred lines studied (Supplemental Figure S1). In summary, our data support a role for bulliform cells in the leaf rolling response, and suggest that rolling is facilitated by more closely spaced and narrower bulliform strips.

**Figure 1.**
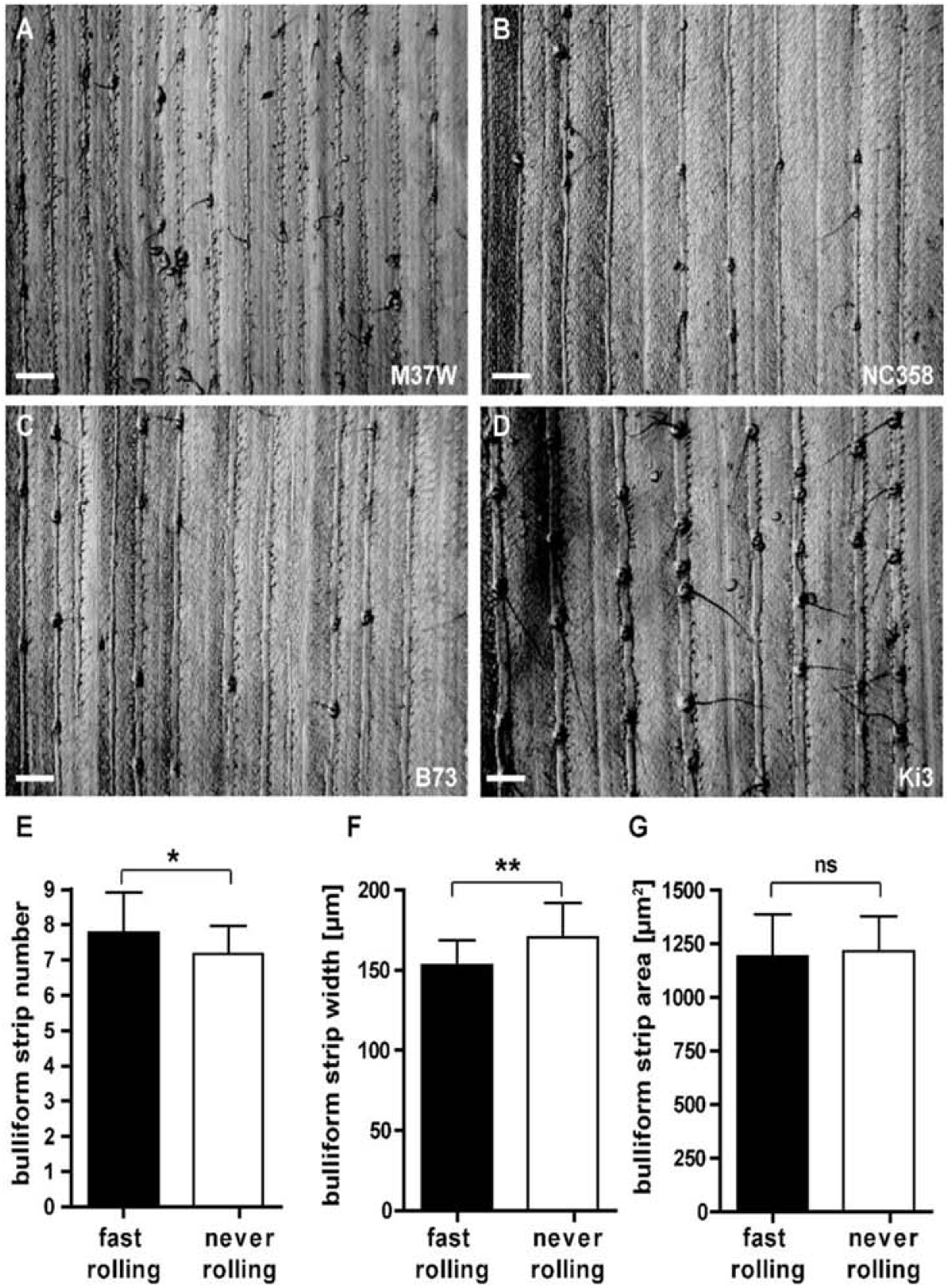
Bulliform patterning correlates with leaf rolling speed. A-D) Grayscale images of leaf epidermal glue-impressions from four maize inbred lines showing extreme bulliform cell patterning phenotypes: M37W and NC358 with narrow bulliform strips, B73 and Ki3 with wide strips. Scale bar = 500 µm. E-G) Comparison of bulliform strip number per field of view with standard size (E), strip width (F) and calculated bulliform strip area (number x width) (G) in 25 fast rolling maize inbreds (rolled within 90 minutes of dehydration; leaf rolling was assessed during a detached leaf dehydration assay in the dark in controlled conditions of 20-22 °C and 55-65 % humidity) and 41 never rolling inbreds (not rolled within 270 minutes of dehydration). Values are given as means ± standard deviation (SD), statistical analysis used two-tailed unpaired Student’s *t*-test, with *P < 0.05, **P < 0.01.

### Bulliform cells display differential shrinkage upon dehydration

Disproportionate shrinkage of bulliform cells, located only on the adaxial side of the leaf, during leaf dehydration is thought to create a hinge-like effect promoting leaf rolling. However, we could not find published experimental evidence confirming this hypothesis. Indeed, it is technically challenging to investigate this, since conventional methods permitting visualization of plant tissues at the cellular level have the potential to cause cell shrinkage in their own right (e.g. fixation and dehydration prior to embedding in a sectioning medium), or reverse cell shrinkage (e.g. if freshly cut hand sections of dehydrated tissue are mounted in aqueous medium under a cover slip). To overcome these problems, a newly-established cryo-confocal imaging method was employed. After 4 hours of dehydration of intact (detached) adult leaves, or no dehydration, tissue fragments were shock-frozen in optimal cutting temperature compound (OCT) and cross-sectioned in a cryo-microtome. Autofluorescence of tissue cross sections exposed at the block face was then imaged in a custom-built liquid N_2_-cooled chamber mounted on a confocal microscope (Figure 2). In comparison with cross sections of fully hydrated control leaves (Figure 2A), pavement cells as well as bulliform cells (red circles) were smaller in dehydrated (rolled) leaves (Figure 2B). Bulliform cell shrank more than pavement cells (Figure 2C,D), but no difference in shrinkage was observed comparing adaxial and abaxial pavement cells.

**Figure 2.**
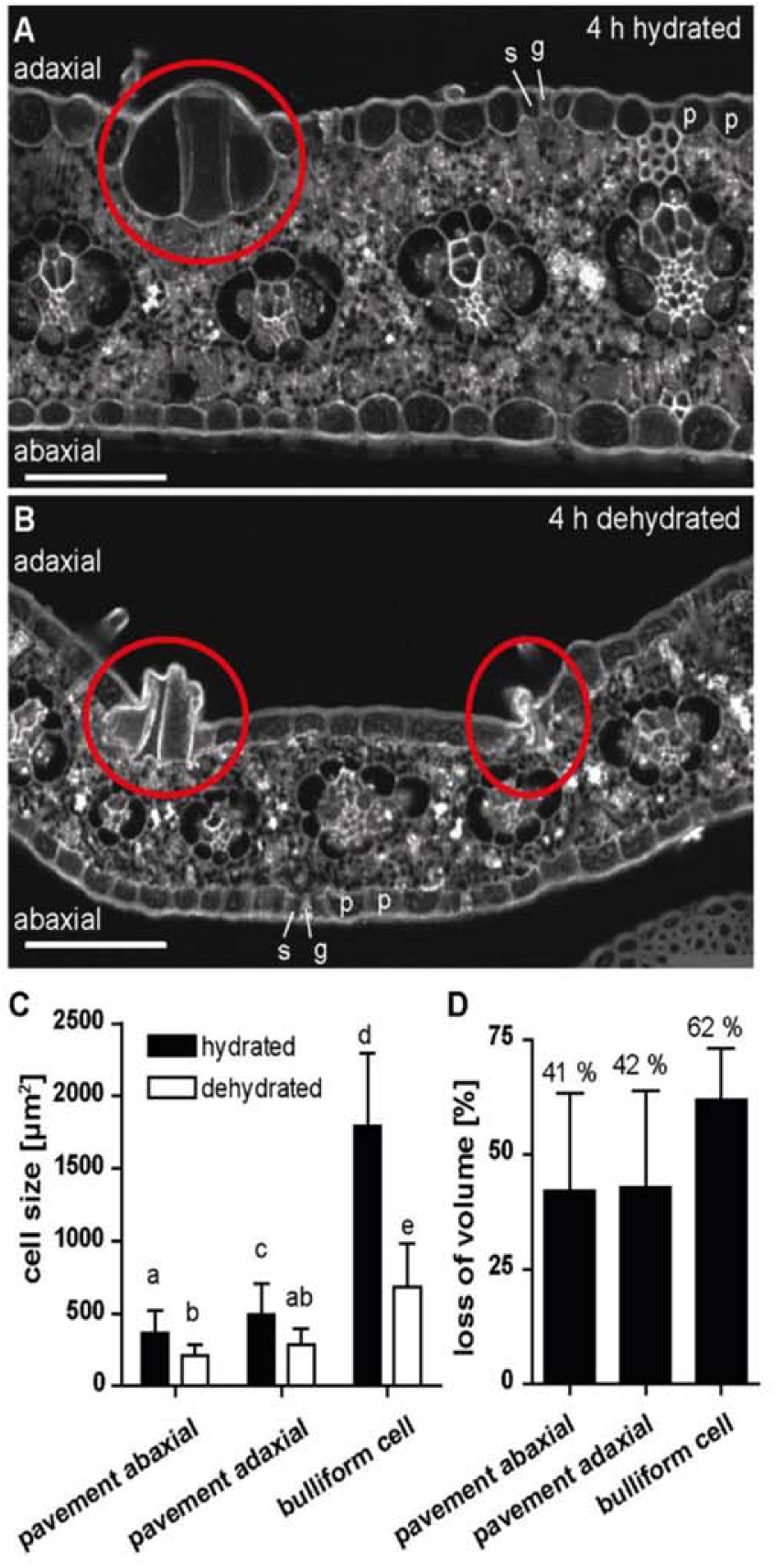
Bulliform cells show increased shrinkage upon dehydration. A-B) Cross sections of (de-)hydrated maize leaf tissue after 4 hours in the dark. Tissue was shock-frozen after 4 hrs without fixation, and autofluorescence was detected by *in situ*-cryo-imaging as described in Methods. p = pavement cell, g = guard cell, s = subsidiary cell, red circle = group of bulliform cells. Scale bar = 100 µm. C) Cell sizes (cross-sectional areas) of the indicated epidermal cell types, analyzed by ImageJ. D) Quantification of cell shrinkage of different cell types upon dehydration as loss of volume in percentage. Shrunken bulliform cells were counted only if their outlines could be seen as in the group on the left in B (i.e. severely shrunken bulliform groups such as the one on the right in B were not counted). Values given as means ± SD (n = 88 - 194 cells per cell type). Statistical analysis used 1-way ANOVA, means with the same letter are not significantly different from each other; Tukey’s post-test, P < 0.05.

To investigate whether differential shrinkage of bulliform cells is due to water loss to the atmosphere (i.e. via evaporation across the bulliform cuticle) or to movement of water into neighboring cells, pavement cell volumes were examined as a function of proximity to bulliform cells and vice versa (Supplemental Figure S2). Bulliform cells lost the same volume upon dehydration, independent of their location in the center of a BC cell strip (#1) or at a peripheral position adjacent to a pavement cell (#2). Moreover, no consistent difference in shrinkage upon dehydration could be observed for pavement cells at different positions relative to BCs (positions #3, #4 and #5, Supplemental Figure S2). Sizes of cells in internal tissue layers could not be determined with the images at hand, so we were not able to investigate the possibility of water transport from bulliform cells into underlying mesophyll cells. Taken together, our results show for the first time *in situ* that dehydration of a leaf leads to increased bulliform cell shrinkage relative to neighboring pavement cells. Analysis of relative cell volumes supports the possibility that water is lost differentially via evaporation across the BC cuticle rather than via transport into neighboring cells.

### Pavement and bulliform cell cuticles of the adult maize leaf are different

Increased volume loss of bulliform cells upon leaf dehydration suggests the hypothesis that differential permeability of bulliform vs. pavement cell cuticles could lead to a difference in water loss. To investigate this hypothesis, cuticle structure was examined in adult, fully mature maize leaves via transmission electron microscopy (TEM) and confocal microscopy (Figure 3). As previously described, TEM reveals that pavement cells have a thin cuticle with four ultrastructurally-defined zones of distinct osmium staining characteristics (Figure 3A, Bourgault et al., 2020). The outermost, dark-staining layer (white arrowhead) represents epicuticular wax layer, while the innermost, dark-staining layer (black arrowhead) could reflect the pectin-rich wall/cuticle interface described for many other plant species (Jeffree, 2006). Between these two layers, darker- and lighter-staining layers can be observed, classified previously as distinct zones of the cuticle proper, since both layers are missing the polysaccharide fibrils characterizing the cuticular layer. Cuticles of bulliform cells (Figure 3B) are strikingly different from those of pavement cells: they are five-fold thicker (Figure 3C), and exhibit a different organization. The epicuticular wax layer (white arrowhead) of BCs is comparable, but the cell wall/cuticle interface (black arrowhead) is diffuse compared to a pavement cell. Dark-staining fibrils (white arrows) are seen reaching into the cuticle from the cuticle-wall interface, mostly aligned perpendicular to the plane of the cuticle; these are characteristic of a cuticular layer with polysaccharides embedded (Jeffree, 2006; Mazurek et al., 2017). A layer devoid of these fibrils (cuticle proper) can be seen directly underneath the epicuticular wax layer. Therefore, bulliform cell cuticles exhibit a standard cuticle consisting of a cuticular layer, cuticle proper and epicuticular wax layer (Jeffree, 2006). A difference between pavement and bulliform cell cuticles was also apparent in leaf cross sections stained with Fluorol yellow 088 (FY) and imaged via confocal microscopy (Figure 3D,E). Whereas pavement cells showed very thin or no FY staining, BCs displayed bright FY staining. A clear, immediate decrease in cuticle thickness can be observed at the boundary between bulliform and pavement cell, where the two cell walls (stained by Calcofluor white (CW)) meet (Figure 3E). In summary, compared to pavement cells, bulliform cells have much thicker cuticle with a prominent cuticular layer.

**Figure 3:**
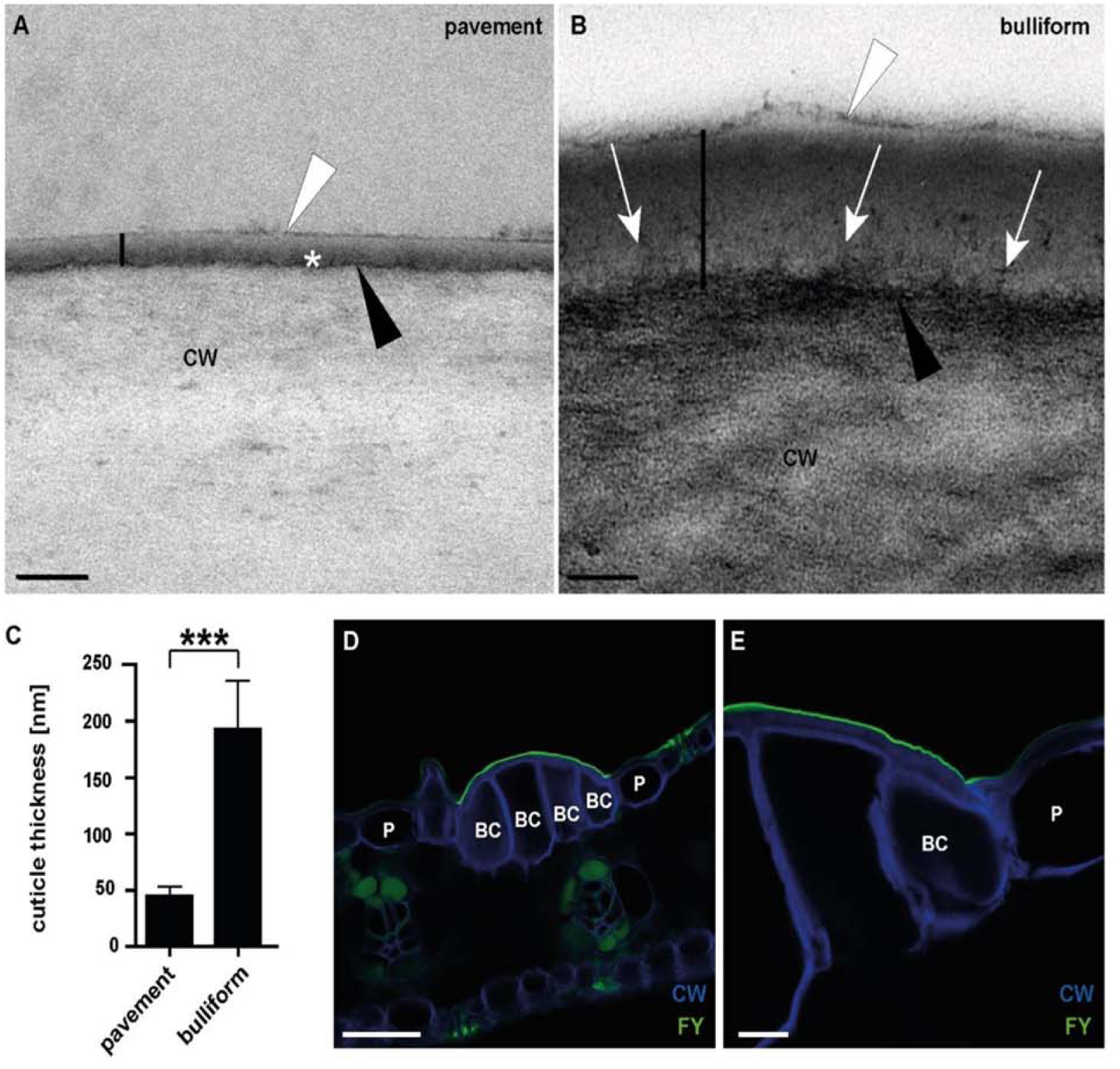
Bulliform and pavement cell cuticles have different thicknesses and ultrastructures. A) Pavement cell cuticle from a fully expanded adult maize leaf, visualized by TEM (vertical black line marks the full extent of the cuticle). Four distinct layers or zones are visible: a thin, darkly stained layer (black arrowhead) at the interface between the cell wall (CW) and cuticle, dark (asterisk) and light zones of the cuticle proper, and a darkly stained epicuticular layer (white arrowhead). B) Bulliform cell cuticle, visualized by TEM (extent marked by vertical black line). The cell wall/cuticle interface (black arrowhead) is diffuse compared to that in pavement cells, and dark-staining fibrils (white arrows) reach from there into the cuticle. White arrowhead points to the epicuticular layer. Scale bar in A) and B) = 100 nm. C) Thickness of different cuticle types, as indicated by the black bars in A) and B). Values given as means ± SD, n = 45 (3 measurements in 3 different images per cuticle type of 5 biological replicates). Statistical analysis used two-tailed unpaired Student’s *t*-test, with ***P < 0.001. D-E) Fluorol yellow staining of leaf cross sections confirms a thicker cuticle over bulliform cells than over the neighboring pavement cells. FY = Fluorol Yellow (lipid stain), CW = Calcofluor White (cell wall counter stain). Scale bar in D) = 50 nm, in E) = 10 nm.

### Bulliform-enriched tissue shows increased epidermal water loss upon dehydration

To investigate the functional significance of the observed differences between BC and pavement cell cuticles, we examined the dehydration response of mutants with bulliform-enriched epidermal tissue. Three different maize mutant lines with previously described epidermal aberrations on adult leaves were identified and analyzed. The *warty2* (*wty2*) mutant is defect in a Tyrosine kinase (Luo et al., 2013), and shows, similar to *wty1,* disordered cell expansion in the leaf blade producing bulliform cell-like epidermal cells forming wart-like textures on both sides of the leaf (Figure 4A,B) (Reynolds et al., 1998; Sylvester and Smith, 2009). Mutants homozygous for a weak allele of *defective kernel1* called *dek1-Dooner* (*dek1-D*), a mutation in a gene encoding a plasma-membrane protein with 21 transmembrane domains and a calpain protease domain, display an increased frequency of bulliform-like cells on both abaxial and adaxial surfaces (Figure 4C) (Becraft et al., 2002). The *extra cell layers1* (*Xcl1*) mutant, with a semi-dominant mutation in an unknown gene, causes extra cell layers with epidermal characteristics similar to bulliform cells on both sides of the leaf blade (Figure 4D) (Kessler et al., 2002). Bulliform or bulliform-like cell identity of the abnormal epidermal cells in the three mutants was confirmed via Fluorol Yellow (FY) staining (Figure 4F-I). In *wty2* and *dek1-D* mutants, enlarged epidermal cells had FY staining characteristics of bulliform cells (Figure 4G,H). In *Xcl1*, epidermal cells overlying the extra (enlarged) cells on both adaxial and abaxial surfaces showed increased FY staining (Figure 4I). Quantification of the cells with increased FY staining revealed an increased proportion of bulliform-like cells in the epidermis of all three mutants (Figure 4E). TEM was conducted to further investigate the cuticles of aberrant, bulliform-like cells in *wty2* and *Xcl* mutants (the *dek1-D* mutant, whose comparably low coverage of abnormal epidermal cells did not allow for a clear identification of these cells in the high magnification but low-throughput setting of TEM, was omitted from this analysis). Aberrant epidermal cells in both *wty2* and *Xcl1* mutants showed ultrastructural characteristics of wild-type bulliform cell cuticles (Figure 5).

**Figure 4:**
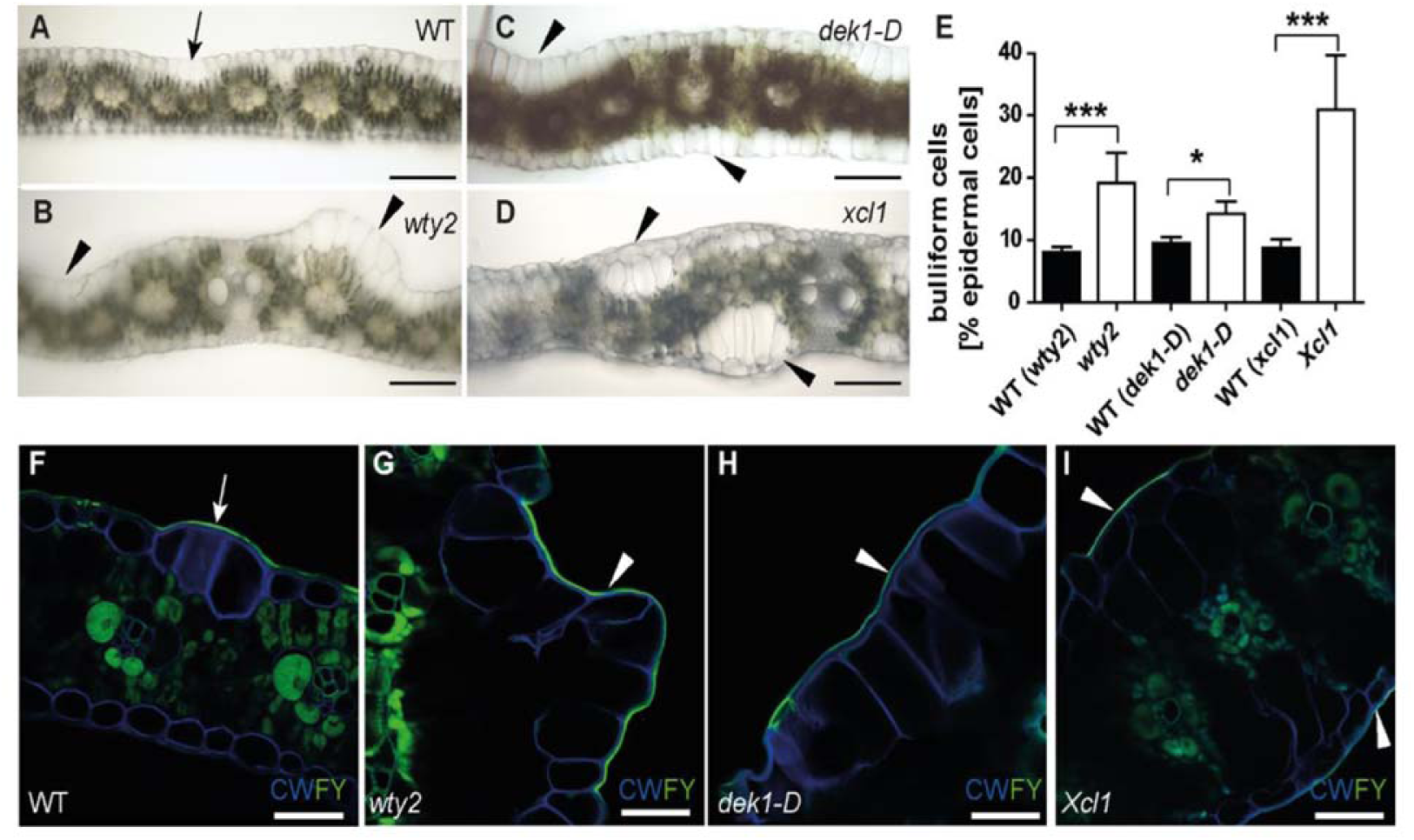
Three different epidermal mutants have increased bulliform cell coverage. A-D) Bright field images of hand-sectioned adult leaves from the indicated genotypes depict previously reported aberrations in epidermal cell types. Arrow = wild-type-like bulliform strip (2 cells wide), arrowhead = areas with abnormal bulliform-like cells. Scale bar = 200 µm. E) Quantification of bulliform cell number in three bulliform mutants as percentage of all epidermal cells, calculated using (F-I) Fluorol Yellow stained tissue cross section images. FY = Fluorol Yellow (lipid stain; green), CW = Calcofluor White (cell wall counter stain; blue). White arrow = FY staining of wild-type-like bulliform strip cuticle (3 cells wide), white arrowhead in G-I) = areas with abnormal BC-like cells displaying increased FY staining of the cuticle. Scale bar = 50 µm. Values in E) are given as means ± SD (n = 300-380 epidermal cells counted, 4 biological replicates per genotype). Statistical analysis used two-tailed unpaired Student’s *t*-test, with *P < 0.05, ***P < 0.001.

**Figure 5:**
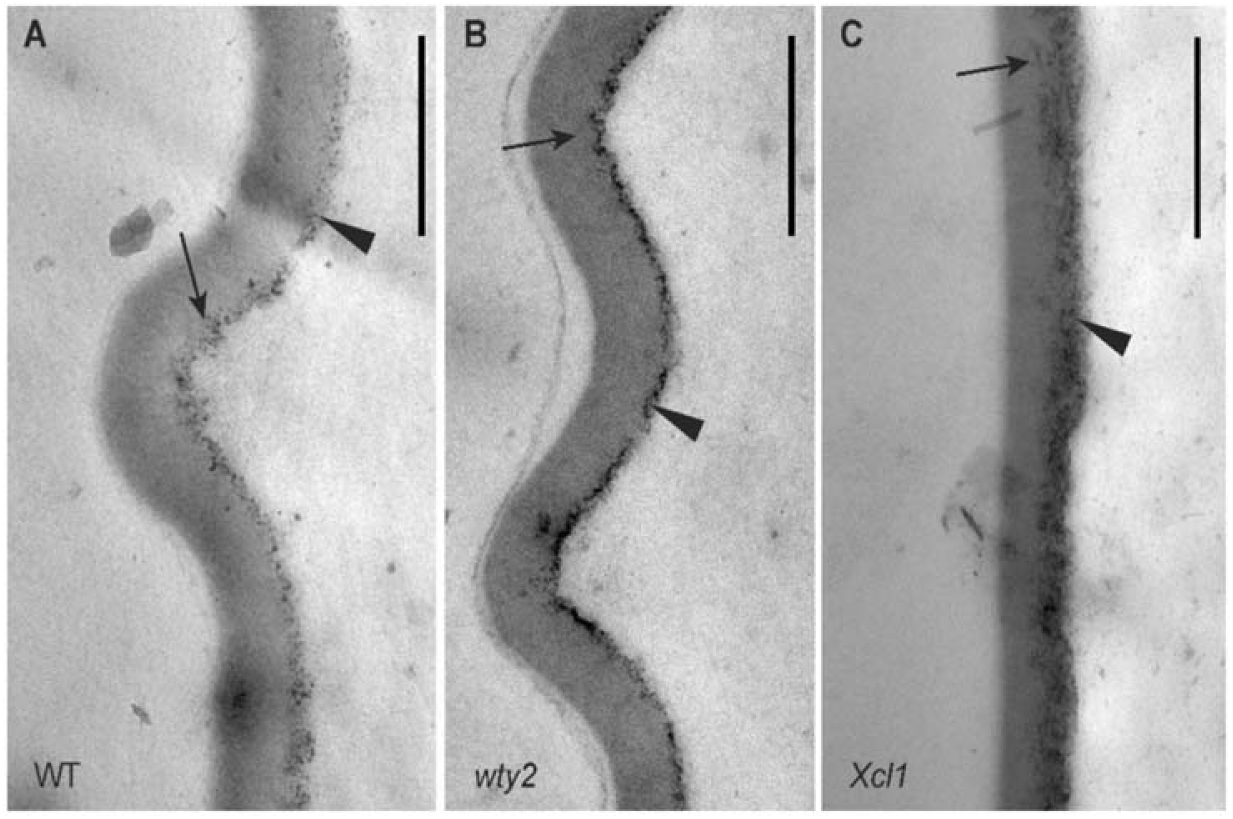
Cuticles of aberrant epidermal cells in bulliform-enriched mutants display BC-like ultrastructure. A-C) Bulliform cell cuticle of wild-type, *wty2*, and *Xcl1* mutant, visualized by TEM. Images in B and C display the outer surface of abnormal bulliform-like cells in the respective mutants, identified by the presence of related abnormal epidermal features of the area (warts in wty2, extra cell layer in *Xcl1*) before acquiring the TEM images. The cell wall/cuticle interface (black arrowhead) is diffuse and dark-staining fibrils (arrows) reach into the cuticle towards the outer surface. Scale bar = 500 nm.

All three epidermal mutants displayed an increased number of bulliform-like cells on their surface, making them adequate tools to further investigate the role of bulliform cells and their cuticles in dehydration (Figure 6). This was assessed by measuring epidermal water loss, also called cuticular conductance (g_c_), of detached leaves in the dark to minimize stomatal water loss (Ristic and Jenks, 2002; Lin et al., 2019). Mature adult leaves of all three bulliform-enriched mutants showed a significantly increased g_c_ compared to their wild-type siblings (Figure 6A), supporting the hypothesis that the bulliform cuticle is more water permeable. To further investigate this hypothesis, a similar dehydration experiment compared g_c_ of adaxial leaf surfaces of adult wild-type leaves (containing bulliform cells), to that of bulliform-free abaxial surfaces. This was achieved by covering one or the other surface with petroleum jelly to prevent water loss from the covered side of the leaf (Fig 6B). Increased dehydration of the bulliform-containing, adaxial side of the leaf was observed compared to the abaxial, bulliform cell-free side, while the control without petroleum jelly constituted the full cuticular conductance. In conclusion, complementary experiments investigating the relationship between bulliform distribution and g_c_ support the hypothesis that bulliform cell cuticles are more permeable to water in spite of their greater thickness, providing a possible mechanism to explain their differential shrinkage, assisting the rolling of grass leaves upon dehydration.

**Figure 6:**
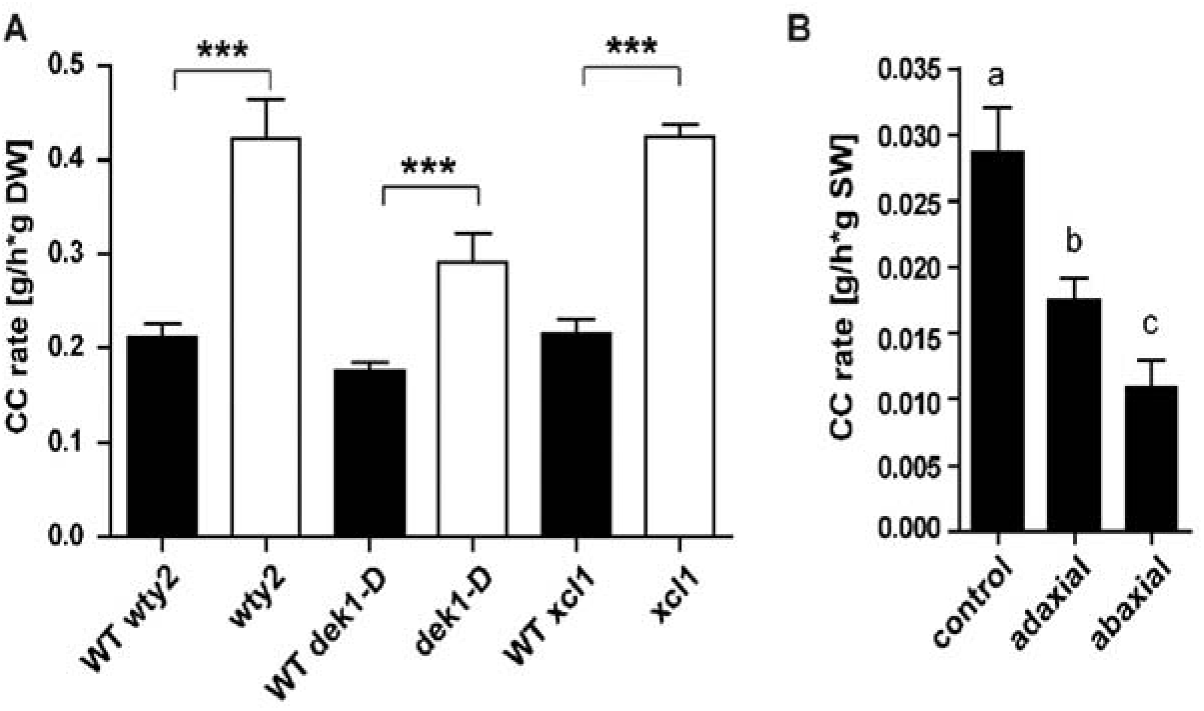
Bulliform-enriched tissues have increased cuticular conductance rates. Cuticular conductance, representing rates of water loss across the cuticle in the dark when stomata are closed, was measured during dehydration of detached leaves (20-22 °C, 55-65 % humidity). A) Cuticular conductance for three BC-enriched mutants *wty2*, *dek1-D*, and *Xcl1* and corresponding wild-types). CC rate is calculated as water loss (g) per hour per g dry weight (DW). Values are given as means ± SD (n = 3-5 biological replicates per genotype). Statistical analysis used two-tailed unpaired Student’s *t*-test, with ***P < 0.001. B) Cuticular conductance rate of adaxial and abaxial leaf surfaces (one-sided dehydration was achieved by covering one side of the leaf with petroleum jelly) compared to full CC rate, calculated as water loss (g) per hour per g starting weight (SW). Values are given as means ± SD (n = 5-6 biological replicates per surface). Statistical analysis used 1-way ANOVA, means with the same letter are not significantly different from each other; Tukey’s post-test, P < 0.05.

### Bulliform cell cuticle nanoridges are not the main driver of increased dehydration

Bulliform cells show a reticulate pattern of cuticle nanoridges on their surfaces (Becraft et al., 2002), increasing cuticular surface area relative to overall cell surface area, and providing a possible mechanism to increase the rate of dehydration of bulliform cells relative to other epidermal cell types. To address the question of whether the observed increase in water loss from bulliform cells and bulliform-enriched tissues could be explained by this mechanism, high resolution surface imaging of leaf glue impressions was performed with a Keyence VHX-6000 digital microscope system (Figure 7). Bulliform cells in wild-type leaves (white arrowheads) were organized in strips of 3-5 cells, and their cuticles displayed nanoridges aligned with the proximodistal axis of the leaf, often appearing to span cell-to-cell boundaries (white arrow) (Figure 7A-C). Adjacent pavement cells (black arrow) lacked these nanoridges. *wty2* mutants displayed normal bulliform strips (white arrowhead) as well as abnormal, epidermal cell bumps which, upon closer inspection, revealed an even denser than normal pattern of cuticular nanoridges (Figure 7D-F). *dek1-D* mutants usually had wider than normal bulliform strips, but cells in these strips varied with respect to cuticular nanoridges: cells in the center had nanoridges, whereas those towards the outer edges had few or no nanoridges (Figure 7G-I). Interestingly, in the last BC mutant, *Xcl1*, abnormal bulliform-like cells displayed no cuticular nanoridges whatsoever (Figure 7J-L). These differences are not due to variations in leaf water content, as all leaves were fully turgid at the time glue impressions were made. Imaging of abaxial leaf impressions of some of the mutants (Supplemental Figure S3) revealed bulliform-like cells on these usually bulliform-free surfaces, where for *dek1-D* nanoridges could be observed in only some areas, while others displayed bulliform-like cells with a smooth surface (Supplemental Figure S3A,B). As for the adaxial surface, no nanoridges could be detected on the abaxial sides of *Xcl1* mutant leaves (Supplemental Figure S3C,D). In conclusion, increased water loss in the three analyzed bulliform-enriched mutants cannot be caused solely by surface area increase due to the presence of cuticle nanoridges on their excess bulliform-like cells, since not all of the mutants displayed these cuticle features despite a higher cuticular conductance rate.

**Figure 7:**
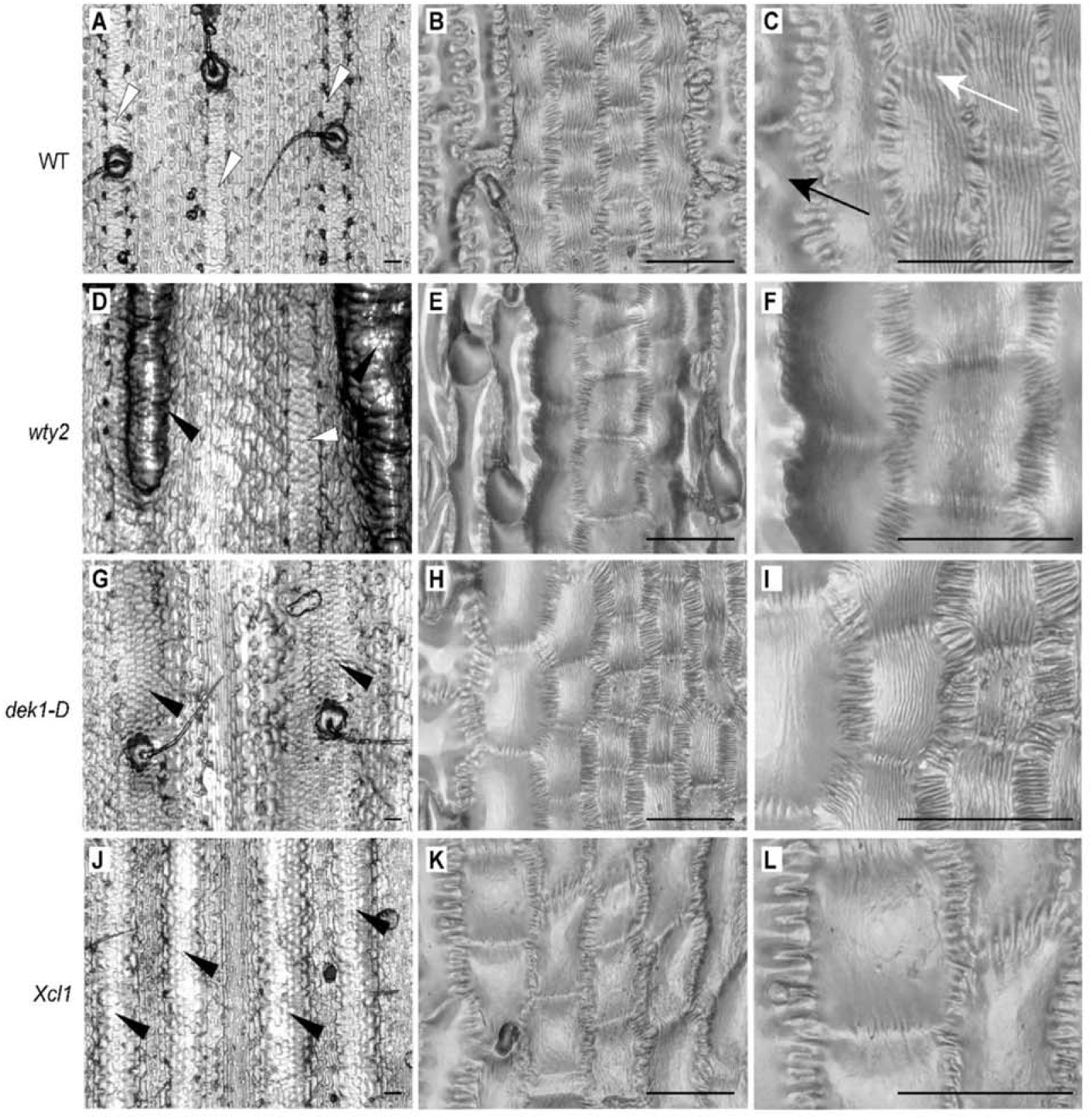
Bulliform-like cells in bulliform enriched mutants do not necessarily have cuticle nanoridges. Epidermal glue impressions of wild-type (A-C) and three bulliform-enriched mutants *wty2* (D-F), *dek1-D* (G-I), and *Xcl1* (J-L) at different magnifications. White arrowheads in A and D point to wild-type-like bulliform strips, while black arrowheads in D, G and J mark abnormal bulliform strips. White arrow depicts nanoridges spanning cell-to-cell boundaries, black arrow indicates a pavement cell without cuticle nanoridges. In the middle and right columns, higher magnification views of bulliform-like cells in each genotype show their surface features in more detail. Note the absence of nanoridges on bulliform-like cells in *Xcl1* mutants. Scale bar in A,D,G,J = 100 µm, in all others 50 µm.

### Bulliform-enriched cuticles have a unique biochemical composition with major differences in cutin

In an effort to identify unique and possibly functionally significant components of bulliform cell cuticles, we sought to biochemically characterize them. Since no method was available to physically separate bulliform from pavement cells on the scale needed to biochemically analyze their cuticles directly, two complementary approaches were taken to compare these cuticle types indirectly with respect to both wax and cutin monomer composition: (1) adaxial (bulliform-containing) and abaxial (bulliform-free) cuticles were compared (see methods for information on how this was achieved), and (2) leaf cuticles of the bulliform-overproducing mutants (*wty2*, *dek1-D,* and *Xcl1*) were compared to their respective wild-type siblings. We reasoned that compositional changes seen in both comparisons should provide insight into the specific composition of the bulliform cell cuticle.

The total lipid polyester (cutin) monomer load of wild-type adaxial cuticles was significantly higher than on the abaxial side (Figure 8A). The relative abundance of individual monomer classes in both adaxial and abaxial cuticles was determined via normalization to the total monomer load on the respective side. Some monomers, including 18:0 FA, several hydroxy FA, and DCAs, were reduced on the adaxial leaf surface (Figure 8A), while one compound, 18:0 9-epoxy-18-OH, showed a major accumulation in the adaxial surface containing BCs. All three bulliform-enriched mutant cuticles (Figure 8B) also had a higher total cutin monomer load compared to wild-type, thus we compared their relative lipid polyester monomer composition. Most monomers found to be different in the adaxial/abaxial comparison did not overlap or overlapped only partially with the differential abundances detected in the bulliform mutant analysis. However, the polar cutin component that is highly enriched in the BC-containing adaxial wild-type cuticles, 18:0 9-epoxy-18-OH, was also significantly increased in all three bulliform-enriched mutant cuticles. These findings point to 18:0 9-epoxy-18-OH as a cutin monomer that is enriched in bulliform cell cuticles.

**Figure 8:**
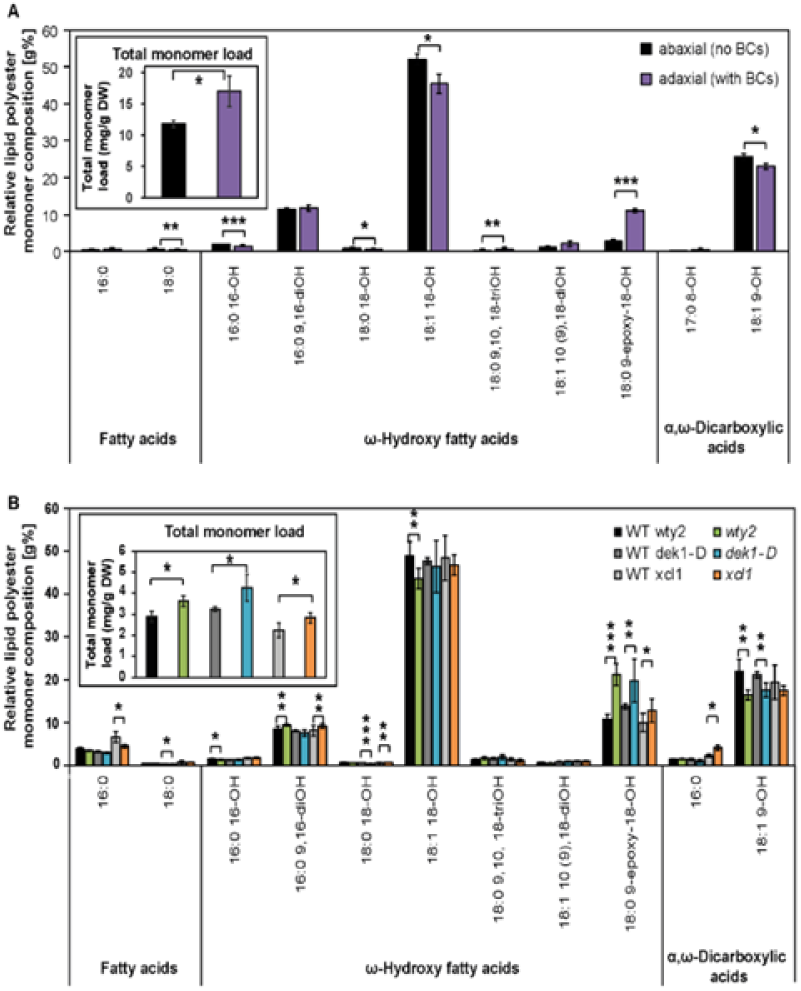
Bulliform-enriched cuticles have a unique biochemical composition with major differences in cutin. A) Representative profile of cutin monomer composition of abaxial (BC-free) and adaxial (BC-containing) adult maize leaf surfaces, extracted and depolymerized from epidermal peels after enzymatic digestion, and measured via GC-MS. Monomer content of single compounds was normalized to overall cutin monomer load (inset). B) Representative profile of cutin monomer composition of bulliform-enriched mutants after whole-tissue extraction and depolymerization, measured via GC-MS. Monomer content of single compounds was normalized to overall cutin monomer load (inset). Values are given as means ± SD (n = 4 biological replicates per surface/genotype). Statistical analysis used two-tailed unpaired Student’s *t*-test, with *P < 0.05, **P < 0.01, and ***P < 0.001.

Cuticular wax analysis of bulliform-enriched tissues created a more complex picture (Figure 9). Total wax load on the adaxial (bulliform-containing) side of wild-type leaves was increased (Figure 9A), but decreased in two out of three of the bulliform-enriched mutants (Figure 9C). Relative amounts of individual wax types are presented in Figures 9B and 9D following normalization to the total wax load. A significant increase in hydrocarbons (alkanes and alkenes) was detected on the BC-containing adaxial side of wild-type leaves (Figure 9B), and hydrocarbons were also enriched in two of the three bulliform-enriched mutants (Figure 9D). The mutant where hydrocarbon enrichment was not observed (*dek1-D*) shows the weakest bulliform enrichment phenotype (Figure 4I), and could have failed to show the hydrocarbon enrichment seen in the other two mutants for this reason. Thus, most of the data support the possibility of hydrocarbon enrichment in bulliform cell cuticular waxes. No other wax classes showed relative abundance changes that were in agreement between adaxial/abaxial comparisons and bulliform mutants vs. wild-type comparisons. While all three mutants showed increased free fatty acid content (Figure 9D), this difference was not seen in the adaxial/abaxial comparison (Figure 9B). Fatty alcohols and wax esters were decreased in two of the three mutants, but this difference was also not observed in the adaxial/abaxial analysis. Aldehydes, which were found to be decreased in adaxial (bulliform-containing) surface waxes, did not show a difference in the bulliform mutant comparisons. Analysis of single compounds within each wax class (Supplemental Figure S4) also did not clearly point to specific wax molecules as specific to or enriched in BC cuticles. Thus, apart from evidence for hydrocarbon enrichment, we found no clear indication from these analyses of enrichment or depletion in individual wax classes or molecules in bulliform cuticles.

**Figure 9:**
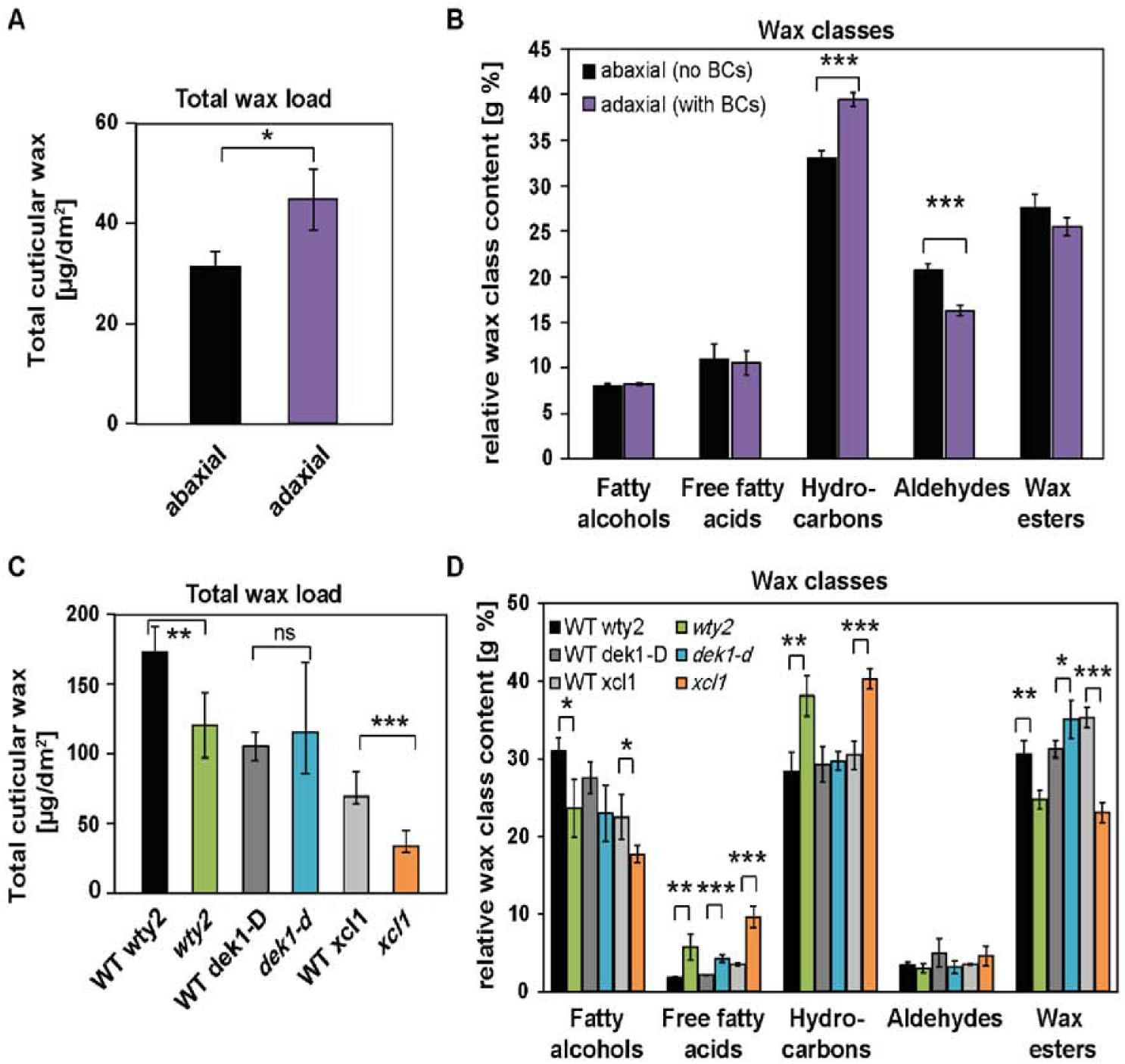
Wax profiles of bulliform-enriched cuticles are diverse. A) Total wax load of abaxial (BC-free) and adaxial (BC-containing) leaf surfaces, chloroform-extracted from one leaf surface or the other, and measured by GC-MS. B) Relative content of wax classes in adaxial and abaxial tissues after normalization to overall wax load. C) Total wax load of bulliform-enriched mutants after chloroform extraction of both leaf surfaces. D) Content of wax classes in bulliform-enriched mutants was normalized to overall wax load. Values are given as means ± SD (n = 4 biological replicates per surface/genotype). Statistical analysis used two-tailed unpaired Student’s *t*-test, with *P < 0.05, **P < 0.01, and ***P < 0.001.

### Transcriptome analysis suggests a role of ferulate in BC cuticle maturation

In order to identify genes whose functions underlie unique features of bulliform cell cuticles, gene expression analysis of the three bulliform mutants was conducted. The data were analyzed to search for genes differentially regulated in the same direction during cuticle maturation in all three mutants compared to their wild-type siblings. To this end, the previously characterized zone of cuticle maturation in developing adult leaves (from 10-30% of the length of a partially expanded leaf #8; Bourgault et al., 2020) was harvested from mutants and corresponding wild-types, and analyzed via RNAseq (Supplemental Figure S5). The three mutants displayed a varying degree of differential gene regulation, with 4833 differentially expressed genes (DEG) for *wty2* in the analyzed zone (Supplemental Figure S5A), comparably less differential gene expression for *dek1-D* with only 132 DEG (Supplemental Figure S5B), and 527 DEG detected for *Xcl1* (Supplemental Figure S5C). However, the overlap between the DEGs in all three mutants was minimal, with only 4 genes showing a differential regulation in all three datasets (Supplemental Figure S5D, Supplemental Table S2), and only two of these genes deviating from wild-type values in the same direction in all three mutants (Supplemental Figure S5E). One of these genes, *Zm00001d008957*, which showed increased expression in all three BC mutants, encodes a putative indole-3-acetic acid-amido synthetase and is annotated as *aas10* (auxin amido synthetase10). BLAST analysis identified AtJAR1, an enzyme responsible for the last step of the biosynthesis of bioactive jasmonic acid (JA), JA-Isoleucine (citation), as its closest relative in *Arabidopsis* (Supplemental Table S3). Other Arabidopsis proteins with high amino-acid sequence identity to AAS10 mostly belong to the auxin-responsive GH3 protein family. The other gene differentially regulated in the same direction in all three BC mutants (reduced expression) is *Zm00001d050455*, which is not functionally annotated in the maize genome but its closest relative in Arabidopsis encodes a hydroxy-cinnamoyl-CoA shikimate/quinate hydroxy-cinnamoyl transferase in Arabidopsis (HCT (Hoffmann et al., 2004), 37% identical at the protein level) (Supplemental Table S4). Interestingly, a slightly less similar gene (28% identity at the protein level, Supplemental Table S4) encodes *DEFICIENT IN CUTIN FERULATE* (*DCF*) in Arabidopsis. Defects in this gene lead to an almost complete absence of ferulate in the cutin fraction of rosette leaf cuticles (Rautengarten et al., 2012). A phylogenetic analysis of selected BAHD family acyltransferases from maize and other organisms (Supplemental Fig. S6, after Molina and Kosma (2015)) positioned the maize candidate Zm00001d050455 (light blue) with several other designated maize “HCT” proteins in a distinct subclade within the HCT clade (dark blue), which might have evolved different functions than HCT. This led us to investigate if the reduced expression of *Zm00001d050455* in bulliform-enriched tissue indeed has an influence on HCA content specifically in the BC cuticle by using our previously described dataset. In the comparison of wild-type adaxial vs. abaxial cutin, isolated from epidermal peels, decrease in HCAs could be observed on the adaxial (bulliform-containing) side (Figure 10A), which was solely due to reduced ferulate content (Figure 10B). Analysis of relative HCA content revealed an increase of coumarate and decrease of ferulate in the adaxial cuticle (Figure 10C). This suggests that BC cuticles are indeed reduced in ferulate, supporting the hypothesis that the putative HCA biosynthetic gene *Zm00001d050455* promotes ferulate incorporation into the polyester and is downregulated in BCs relative to other epidermal cells (Supplemental Figure S5) leading to reduced ferulate content of BC cuticles. This analysis could not be readily extended to BC-enriched mutants because cutin analysis conducted on these mutants utilized whole leaves (not epidermal peels), thereby including internal tissue. Previous work showed that that most HCAs in samples prepared this way are of non-epidermal origin (Bourgault et al., 2020) so they are not informative regarding cuticular HCA content. In summary, analysis of gene expression in BC-enriched mutants suggests roles for hormone and HCA biosynthesis in BC differentiation. These results combined with analysis of HCA content in adaxial vs. abaxial cuticles suggest that ferulate biosynthesis is downregulated during cuticle maturation to achieve reduced ferulate content in BC cuticles relative to other epidermal cell types.

**Figure 10:**
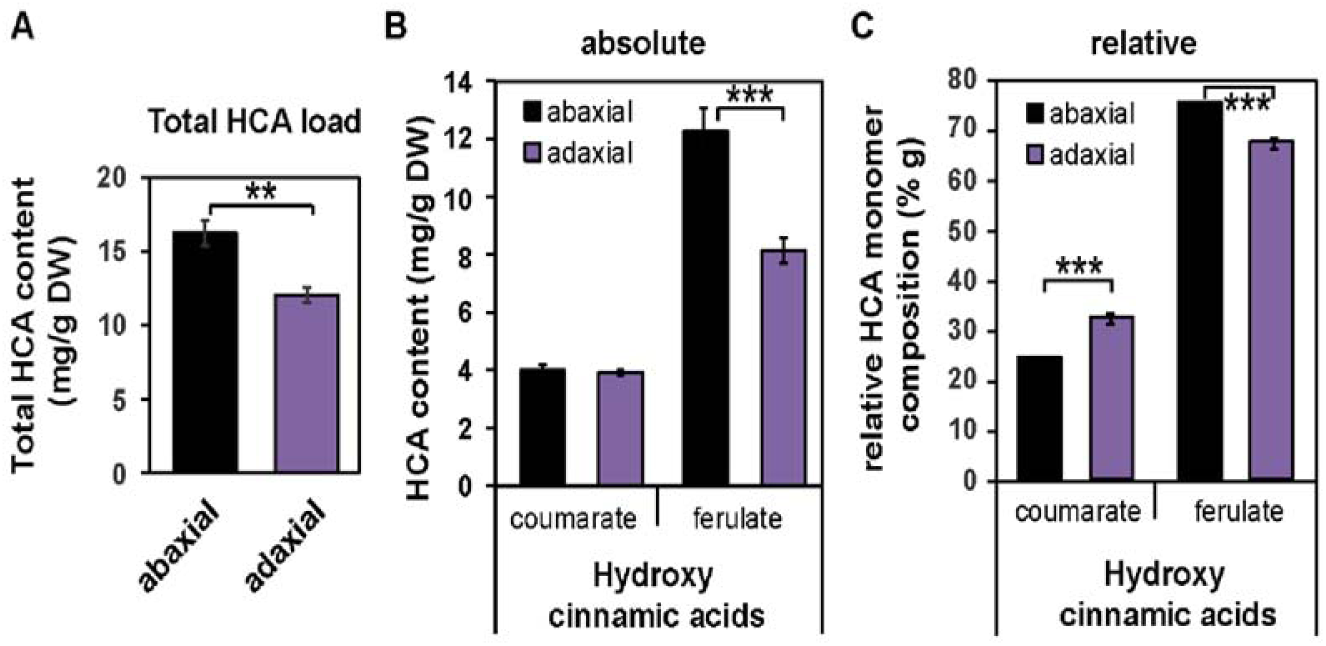
Ferulate is depleted in the adaxial (bulliform-containing) cuticle. A) Total hydroxycinnamic acid (HCA) content of abaxial and adaxial leaf surfaces, extracted and depolymerized from epidermal peels after enzymatic digestion, and measured by GC-MS. B) Absolute and C) Relative content of coumarate and ferulate in abaxial vs. adaxial tissues. Values are given as means ± SD (n = 4 biological replicates per surface). Statistical analysis used two-tailed unpaired Student’s *t*-test, with **P < 0.01 and ***P < 0.001.

## Discussion

Variability in cuticle composition and structure is found between different plant species, developmental stages and tissue types (Jeffree, 2006; Jetter et al., 2006). But even within a tissue there are different cell types carrying out specific functions, which might require different surface properties and consequently cuticle types to support these functions. As one of the first examples of a structure-function relationship of a cell type-specific cuticle, this study set out to investigate BC cuticles in maize and their putative functional role in leaf rolling. Natural variation in maize was used to identify architectural features of bulliform strip distribution associated with leaf rolling speed, emphasizing the importance of bulliform strip architecture for leaf rolling. Ultrastructural and biochemical analyses were carried out to relate BC cuticle features to the unique wax and cutin composition of BCs, pointing to major differences in cutin load and composition, and thus presumably cutin structure in the BC cuticle. Functional analyses revealed increased water loss rates for BCs in dehydrating leaves, probably over the cuticle surface. These findings support the hypothesis that BC cuticles are more water permeable than pavement cell cuticles, possibly facilitating the function of bulliform cells in stress-induced leaf rolling of grasses.

### Maize leaf rolling is impacted by variation in bulliform strip patterning

The role of BCs in leaf rolling has been a matter of ongoing debate for decades, and no clear conclusion has been drawn about their functional contribution to this important drought response (Ellis, 1976; Moulia, 2000). Loss of turgor in bulliform cells on the adaxial leaf surface has long been thought to induce rolling, with additional contribution by shrinkage of subepidermal sclerenchyma and mesophyll tissue due to water loss (Redmann, 1985). But rolling can also occur in leaves that lack bulliform cells (Shields, 1951), questioning the necessity of this cell type for the leaf rolling response. The present study attempted to further establish a functional role for BCs in leaf rolling in maize by examining the relationship between bulliform patterning and leaf rolling across a large collection of genetically diverse maize lines. Data on bulliform strip pattern collected for a GWAS of this trait (Qiao et al., 2019) were analyzed in relation to leaf rolling rate data collected for the same plants. Faster rolling speed was positively correlated with bulliform strip frequency and negatively correlated with bulliform strip width, indicating that rolling is facilitated by more closely spaced and narrower bulliform strips. Nevertheless, to our knowledge, no other study has analyzed the impact of BC architectural variation in grasses on leaf rolling. A study on the flag leaf in wheat revealed that a drought-resistant variety, exhibiting faster leaf rolling than a comparable drought-susceptible genotype, had larger BCs, possibly contributing to the faster rolling response, but also other altered features like differences in cuticular composition (Willick et al., 2018). No information about the BC architecture across the leaf was collected in this case.

Our findings add to prior observations that many leaf rolling mutants in rice or maize show alterations in BC size, number or adaxial/abaxial positioning (Nelson et al., 2002; Xu et al., 2018; Gao et al., 2019). Leaf architecture in general seems to be important, as histological analyses of a collection of 46 adaxially or abaxially rolled mutants in rice showed that changes of number, size, and pattern of bulliform cells, sclerenchyma cells, parenchyma cells, and mesophyll cells as well as vascular bundles all could cause altered leaf rolling (Zou et al., 2014). However, these mutations usually lead to a constantly rolled leaf status rather than alterations in the inducibility or speed of rolling upon drought or heat stress. In general, our data support the conclusion that bulliform strip architecture and distribution across the leaf play a role in regulation of leaf rolling. Although a microscopic phenotype, bulliform strip patterning could represent an important agronomic trait with consequences on macroscopic phenotypes such as plant architecture and drought resistance.

### BC cuticles of the adult maize leaf are structurally and compositionally unique

In the adult maize leaf, pavement cells, BCs, stomatal guard, and subsidiary cells all show different cuticle ultrastructure (Bourgault et al., 2020). The present study examined the ultrastructure of BC cuticles in detail. BCs exhibit a roughly 4-fold thicker cuticle with a prominent cuticular layer, which was not evident in pavement cell cuticles. This cuticular layer is ultrastructurally defined by the presence of osmiophilic fibrils oriented perpendicular to the plane of the cuticle, likely to be polysaccharides (Jeffree, 2006; Mazurek et al., 2017). This likens the BC cuticle more to the classic three-layered cuticle model than the pavement cell cuticle of maize leaves, which lack a well-defined cuticular layer but have two distinct layers of the cuticle proper (Bourgault et al., 2020).

This study also investigated the composition of the BC cuticle. The thickness and ultrastructure of pavement cell cuticles in adult maize leaves are indistinguishable on adaxial and abaxial surfaces (Bourgault et al., 2020). Thus, differences in cuticle composition between the two surfaces are likely due primarily to the presence of BCs and other cell types (e.g. hairs) present only on the adaxial side. To exclude contributions of other cell type-specific cuticles, a complementary analysis of bulliform-enriched tissue was undertaken by comparing three different bulliform-enriched mutants (*wty2*, *dek1-D* and *Xcl1*) to their respective wild-type siblings. Differences observed in both comparisons (bulliform mutants vs. wild-type, and adaxial vs. abaxial) very likely depict a true compositional difference between BC and pavement cell cuticles, and might indicate important functional components of this special cuticle type.

An overall increase of cutin (but not wax) load was found in bulliform-containing or -enriched tissue in all the comparisons. This increase is consistent with the dramatically increased thickness of the BC cuticle, an increase that is mostly due to the presence of a (presumed cutin-containing) cuticular layer not present in pavement cell cuticles. Moreover, all comparisons agreed in identifying the cutin monomer 9,10-epoxy-18-hydroxyoctadecanoic acid as being highly enriched in bulliform cuticles. Together, our findings identify cutin load and monomer composition as the main differences between BC and pavement cell cuticles, potentially changing the physical properties of the cuticle due to different degrees of cross-linking in the polymer scaffold (Fich et al., 2016). Analysis of the petal cuticle in Arabidopsis revealed that cutin biosynthesis is required for the formation of cuticle nanoridges (Li-Beisson et al., 2009; Mazurek et al., 2017), supporting the idea that nanoridges found on the BC cuticle might be present due to different cutin load and/or composition, likely resulting a different polyester structure, compared to pavement cells, which lack nanoridges. The functional significance of 9,10-epoxy-18-hydroxyoctadecanoic acid in the BC cuticle is unclear, but it has previously been implicated in freeze-resistance of cold-hardened rye (Griffith et al., 1985). Interestingly, this monomer was found to be the dominating cutin component in the mature leaf portion of *Clivia miniata*, where its accumulation is thought to indicate that the possible maximum of cross-linking in the cutin fraction had not been achieved (Riederer and Schönherr, 1988). High content of monomers with unused functional groups for cross-linking, like epoxy substituents, can indicate a reduced proportion of actual cross-linking (Riederer and Schönherr, 1988) suggesting a looser cutin scaffold.

The ability to draw definitive conclusions about differences in cuticle composition between BCs and other epidermal cell types is limited by the inability to isolate these cells in sufficiently large quantities for biochemical analysis of cuticles. A promising avenue for future analyses of cuticle specializations in BCs and other epidermal cell types would be the employment of single-cell *in-situ*-imaging techniques, previously shown with e.g. Infrared (IR), FTIR (Fourier transform IR) (Mazurek et al., 2013), Raman scattering spectroscopy techniques (Yu et al., 2008; Weissflog et al., 2010), or compositional analysis of waxes or cutin via matrix-assisted laser desorption/ionization mass spectrometry imaging (MALDI-MSI) (Cha et al., 2009; Veličkovic et al., 2014). Jun and colleagues (2010) achieved a spatial resolution of ∼12 μm in MS-imaging of flower tissue in Arabidopsis, with single pixel profiles demonstrating single-cell-level spatial resolution, allowing for semi-quantification of surface metabolites on single pixels of the flower tissue. Although these techniques require very specialized equipment and expertise, application of these and related methods seem to be able to deliver cell-to-cell resolution of cuticle composition.

Gene expression analysis of BC-enriched mutants yielded fewer candidates regulators of BC differentiation than anticipated, but identified two genes that were differentially expressed in the same direction the cuticle maturation zone of all three mutants compared to wild-type. One of these, *Zm00001d050455,* is a homolog of *Arabidopsis HCT*, which is involved in phenylpropanoid biosynthesis (Hoffmann et al., 2004). Another member of this gene family in Arabidopsis, *AtDCF*, encodes a protein that promotes ferulate esterification into the growing cutin polymer (Rautengarten et al., 2012). The reduced expression of *Zm00001d050455* observed in all 3 BC-enriched mutants could suggest a possible role for this gene in HCA deposition in cuticles. While a function similar to HCT would predict a role in ferulate biosynthesis, a function similar to DCF would suggest reduced incorporation of ferulate into the polyester upon downregulation of the gene in BCs. Phylogenetic analysis of members of the BAHD acyltransferase family positioned the candidate closer to the HCT clade than to proteins related to extracellular lipid biosynthesis like DCF, but the localization in a distinct subclade with several other designated “HCT“ maize proteins might suggest that these genes might not be true homologs of HCT either and could have other distinct functions that affect hydroxycinnamic acid accumulation, maybe even specifically in the cuticle. In any case, comparative analysis of abaxial vs. adaxial cutin composition provided evidence that BC cuticles do indeed have reduced ferulate content compared to pavement cells, but a role for *Zm00001d050455* in establishing this difference remains to be confirmed. In general, the functional role of ferulate in the cuticle remains unclear, but analyses of cuticle permeability in Arabidopsis mutants lacking the *DCF* gene suggest that cutin-bound ferulate does not affect structural and sealing properties of the cuticle (Rautengarten et al., 2012). Just as single cell analysis would provide more definitive insights into cell type differences in cuticle composition, more precise and definitive information about gene expression differences underlying the unique structural and compositional features of BC cuticles could be achieved in future work using single-cell RNA-sequencing strategies after isolating BCs from the cuticle maturation zone by techniques such as laser-capture microdissection or microfluidics (Chen et al., 2019).

### Bulliform-enriched tissue shows increased water loss upon dehydration

Several lines of evidence gathered in our study point to the conclusion that the bulliform cuticle could be more water permeable, leading to higher water loss of bulliform cells upon dehydration: 1) BCs show increased volume loss upon dehydration compared to adjacent pavement cells, as shown in the cryo-confocal analysis of dehydrated tissue *in situ*. Water lost from BCs does not seem to be redistributed into the neighboring epidermal cells, as no decreasing gradient in volume loss of adjacent cells respective to their position to BCs was detected, which would be expected upon additional water entry to these cells from the BCs. Partial direction of water flow from BCs to mesophyll tissue cannot be excluded but could not be quantified with the images at hand. Indeed, Haberlandt and Drummond (1928) state that in rapidly transpiring organs the epidermis loses water to photosynthetic tissue with its higher osmotic pressure. Nevertheless, at least some of the water lost by BCs in this process has to cross the water barrier of the cuticle and exit the tissue, since the overall weight of the leaf is decreasing over time, as seen in our and many other studies using this or similar methods to evaluate cuticular conductance (Kerstiens, 1996; Ristic and Jenks, 2002; Lin et al., 2019). Importantly, all our dehydration experiments, including this imagining experiment, were conducted in the dark to minimize stomatal transpiration, so increased volume loss of BCs upon dehydration suggests increased water loss over the cuticle. 2) Bulliform-enriched tissue shows higher water loss rates in detached leaf drying assays than comparable control tissue. Three different mutants with elevated bulliform cell surface areas showed increased cuticular conductance compared to their wild-type siblings, and wild type leaf adaxial leaf surfaces containing BCs displayed higher cuticular conductance than the abaxial surfaces lacking BCs. While studies of cuticular conductance have investigated this trait in a multitude of plant species (e.g. Table 1 in Kerstiens, 1996), only a few compare the g_c_ of adaxial and abaxial tissues. For example, no difference in cuticular conductance between adaxial and abaxial leaf surfaces could be measured for holm oak (Fernández et al., 2014) or beech (Hoad et al., 1996). While these are dicot species lacking BCs on their adaxial surface, a study in rice also showed increased adaxial cuticular conductance in leaves (Agarie et al., 1998), in agreement with our results of higher water loss of bulliform-containing tissue. Again, also under these circumstances water must be crossing the cuticle surface, since there is overall loss of water from the tissue shown by weight decrease of the leaf over time. These findings lead us to speculate that, indeed, the presence of bulliform cells could drive increased cuticular conductance, possibly facilitating leaf rolling upon dehydration.

Is it likely that the much thicker BC cuticle, with major changes in cutin and less in waxes, is more water permeable than other epidermal cuticle types? While some data connect a thicker cuticle to a lower cuticular water loss rate in maize (Ristic and Jenks, 2002), there is a long line of evidence that cuticle thickness in general cannot be taken as measure for water permeance (Riederer and Schreiber, 2001; Jetter and Riederer, 2016). Cuticle composition rather than thickness seems to be the determining factor for cuticular permeability, which generally is attributed to wax components rather than cutin monomers (Kerstiens, 1996; Buschhaus and Jetter, 2012; Jetter and Riederer, 2016). Indeed, several mutants with altered wax or cutin composition showed higher cuticle permeability despite increased cuticle thickness (Xiao et al., 2004; Kurdyukov et al., 2006; Sadler et al., 2016). The accumulation of epoxy-monomers in the cutin fraction of BCs might be an indication of a less-cross-linked cutin scaffold in these cells, since cuticles high in these monomers with unused functional groups for cross-linking are assumed to show a lower actual degree of cross-linking (Riederer and Schönherr, 1988). This difference in polymer structure might lead to the necessity of increased cuticle thickness to still be able to provide adequate water barrier properties of this specialized cuticle, even if more water permeable than other cell type cuticles. Additionally, existence of a layer with polysaccharide fibrils in the BC cuticle could indicate an aqueous connection for easier water passage through the matrix of cutin and waxes with hydrophilic domains provided by polysaccharides (Fernández et al., 2017), while the cuticular layer and polysaccharide fibrils are absent in pavement cells.

In conclusion, we demonstrate that the maize BCs show higher water loss upon dehydration compared to other epidermal cells. The exact role of BCs and their specialized cuticle in the leaf rolling response of maize has yet to be elucidated, but leaf rolling appears to be facilitated by this thicker, yet likely more water permeable cuticle type unique to BCs. Integration of biochemical, transcriptomic, ultrastructural, and functional data suggest an important role for the cutin matrix in this cuticle type, including the compound 9,10-epoxy-18-hydroxyoctadecanoic acid. Together, our findings advance knowledge of cuticle composition/structure/function relationships, and how cuticle specialization can contribute to cell and organ functions.

## Material and Methods

### Plant material and growth conditions

Maize inbred B73 was used for experiments unless otherwise stated. All mutants analyzed were introgressed into the B73 background. *wty2* seeds were obtained from Prof. Anne Sylvester (University of Wyoming), *dek1-D* seeds from Prof. Phil Becraft (Iowa State University), and *Xcl1* seeds from Prof. Neelima Sinha (UC Davis). Plant materials and experimental field designs for the leaf rolling analysis have been described previously (Lin et al., 2019; Qiao et al., 2019). For histological, biochemical, and functional analyses, plants were grown in 8-inch pots in a glasshouse on the UCSD campus in La Jolla, CA (latitude 32.8856, longitude −117.2297), without supplementary lighting or humidity control, and with temperatures in the range of 18–30°C. All experiments presented focused on fully expanded adult leaves before or during flowering stage, starting with the first fully adult leaf (#8 in B73) or concentrating on the leaf subtending the uppermost ear, or one leaf above or below.

### Cuticular conductance

Cuticular conductance was determined as described previously (Lin et al., 2019). In short, whole adult leaves (3-5 per genotype) were cut 2.5 cm below the ligule and incubated in a dark, well-ventilated room for 2 h at 20-22 °C and 55-65% RH, with cut ends immersed in water for stomatal closure and full hydration (porometer studies established that 2 h was more than sufficient to reach g_min_ indicating stomatal closure; Lin et al., 2019). After removal of excess water on the leaf blades, leaves were hung to dry in the same dark, temperature-and humidity-controlled room. To determine g_c_, wet weight of each leaf was recorded every 45 - 60 min over a time period of 270 - 300 min, for a total of five or six measurements per leaf. Leaf dry weight was acquired after 4 days of incubation at 60 °C in a forced-air oven. Dry weight was shown to be a reasonable approximation of leaf surface area for normalization of g_c_ (Lin et al., 2019), and was used in the calculation of adult leaf cuticular conductance as follows (g_c_): g_c_ (g/h*g) = - b / dry weight, where b (g/h) is the coefficient of the linear regression of leaf wet weight (g) on time (h), and dry weight (g) is an approximation of leaf surface area. In case of petroleum jelly treatment of adaxial or abaxial leaf surfaces, weight loss over time was normalized to starting weight since complete drying of petroleum jelly-treated leaves was not possible.

### Leaf rolling analysis

Leaf rolling was scored on a set of 468 maize inbred lines from the Wisconsin Diversity panel (Hansey et al., 2011), which at the same time was evaluated for genetic variation of bulliform patterning (Qiao et al., 2019) and leaf cuticular conductance (g_c_) of adult maize leaves (Lin et al., 2019). Data on leaf rolling (Supplemental Table S1) was collected during the phenotypic evaluation of g_c_ in 2016 at the Maricopa Agricultural Center, Maricopa, AZ. Leaf rolling was recorded during each weight recording for g_c_ analysis, in 45 minutes intervals at 6 time points (TPs) over a span of 270 minutes, using a visual scale of 0 = not rolled, and 1 = rolled. Our score 1 corresponded to score 5 of fully rolled leaves according to Moulia (1994) (with maize belonging to rolling type 2), while score 0 (not rolled) corresponded to scores 1-4 in Moulia’s scoring scale. The TP when leaves were scored to be rolling for the first time (TP1 corresponding to 45 min of dehydration, TP6 corresponding to 270 min), were extracted for each inbred, while leaves which were scored to be unrolled at TP6 got assigned the hypothetical rolling time point TP7 as the most conservative estimation. Pearson’s correlations between bulliform patterning (number and width of BC strips, Supplemental Table S1, extracted BLUPs for the Maricopa/AZ environment from Qiao et al., 2019) and leaf rolling data was analyzed for 291 inbreds in total (overlap between datasets of 316 lines with data for leaf rolling and dataset of 410 lines with data for bulliform patterning). Lines with extreme rolling behavior - “fast rollers” (rolled within the first 90 minutes of dehydration, 25 lines), and “never rollers” (no rolling observed in the assessed time frame, 41 lines) – were grouped and additionally graphed independently.

### Cryo-microscopy of dehydrated leaves

Adult B73 leaves were cut below the ligule and hung to dry in the dark for 4 h, while control leaves were kept hydrated by submerging the cut end in water. Immediately after, blade tissue from the mid-section of the leaves was cut, submerged in cryomolds containing room-temperature optimal cutting temperature compound (OCT, Sakura Finetek USA) and frozen in liquid nitrogen. The frozen OCT blocks were milled flat using a Leica CM 1950 cyrostat, exposing the leaf tissue remaining in the block at the biologically relevant level while discarding the remnants of the cryosections. The frozen block face with the exposed, intact leaf tissue was transferred to a Nikon A1plus Eclipse Ni-E confocal microscope, equipped with a liquid nitrogen-cooled imaging chamber custom-built by Paul Steinbach (laboratory of Dr. Roger Tsien, UCSD), designed to maintain the frozen samples at ∼40° C, and imaged directly with the confocal microscope with a 4X (NA 2.0) objective. Autofluorescence of plant tissue was detected with excitation at 405 nm and emission detection at 525 nm. Cell size measurements were done with ImageJ (v1.50i, https://imagej.nih.gov/ij/).

### TEM and other Imaging

TEM sample preparation and imaging was done as previously described (Bourgault et al., 2020). Glue impressions of adult maize leaves were collected as described previously (Qiao et al., 2019). For examples of natural variation of bulliform strip patterning in Figure 1A-D, impressions of different inbreds were imaged with a Nikon SMZ-U Stereoscopic Zoom Microscope, using a 0.5 objective lens with a Lumenera InfinityX camera attached. For analysis of cuticular nanoridges, glue impressions of bulliform mutants were imaged with a Keyence VHX-6000 digital microscope, equipped with a VH-ZST lens. Hand cross sections of adult leaf blade material of bulliform mutants were imaged on a Nikon Eclipse E600 Microscope with a Lumenera InfinityX camera attached.

### Fluorol Yellow staining

Tissue samples of the middle section of adult leaves were collected and fixed in Formalin-Acid-Alcohol (ethanol (>90%) 50 %, glacial acetic acid 5 %, formalin (37% formaldehyde) 10%). Fixed samples were infiltrated in gradual increases to 30% sucrose solution, embedded in OCT compound (Sakura Finetek USA) and frozen into cryoblocks. 20 µm block face sections were collected on 1% polyethylenimine (PEI) coated slides, stained with 0.1% calcofluor white (aq., Sigma Aldrich) for five minutes followed by 0.01% Fluorol Yellow (Santa Cruz Biotechnology) in lactic acid solution for 30 minutes, mounted in Vectashield anti-fade mounting medium (VECTOR Laboratories), and sealed underneath a coverslip by nail polish. Images were captured on a Zeiss LSM 880 Confocal with FAST Airyscan using Plan-Apochromat 10X/0.45 M27, Plan-Apochromat 20x/0.8 M27, Plan-Apochromat 63x/1.4 Oil DIC M27 objectives set at 515nm emission/488nm excitation wavelength for Fluorol Yellow and 450nm emission/405nm excitation wavelength for calcofluor white. Collected images were processed through superresolution Airyscan and composite pictures were processed through ImageJ. Cell counts of epidermal cell types were done with ImageJ.

### Analysis of leaf cuticular lipids

Abaxial and adaxial B73 cuticles: Adaxial and abaxial leaf surfaces were used from the portion between 20 and 42 cm of maize B73 partially expanded leaf #8 after removing the midrib. Total waxes were extracted following the method described in Buschhaus et al. (2007) with some modifications. A 25 ml glass tube containing 8 ml chloroform was placed against the leaf surface held to the rim of the tube using mild thumb pressure. The chloroform was shaken against the surface of the leaf for 30 seconds. This was repeated along the entire 22 cm length of the leaf and each leaf-half was used for either abaxial or adaxial extraction. Internal standards were added to the extracts; 1.5 μg of each, n-tetracosane (24:0 alkane), 1-pentadecanol (15:0-OH) and heptadecanoic acid (17:0). Extracts were dried under a nitrogen stream and analyzed as described in Bourgault et al. (2020). To determine cutin monomer composition of adaxial and abaxial leaf surfaces, cuticles were isolated from the same portion of B73 leaves described above, using enzymatic digestion (Bourgault et al., 2020). Isolated cuticles were delipidated and dried under a nitrogen stream. Dry weight was recorded at this stage and 10 μg internal standards were added to all extracts; pentadecanolactone (C15:0 ring) and methyl heptadecanoic acid (17:0). Samples were then depolymerized and the released monomers analyzed by GC-MS as described in Bourgault et al. (2020).

Bulliform mutant cuticles: Total waxes were extracted by chloroform immersion from bulliform-overproducing mutants (*wty2*, *dek1-D*, and *Xcl1*) and their respective wild-type siblings using expanded segments of adult leaves. Tissues remaining after chloroform extraction were delipidated and chemically depolymerized. Both wax extracts and cutin monomers were transformed into their TMSi derivatives and analyzed by GC-MS and GC-FID following the same procedures described in Bourgault et al. (2020).

### RNAseq analysis

Total plant RNA of developing adult leaves (10-30 % of leaf length of the maturing leaf at 50 to 60 cm, where cuticle maturation is most prominent according to Bourgault et al., 2020) was isolated with the RNeasy Plant Mini Kit (Qiagen) according to the manufacturer’s instructions. Library preparation and RNAseq were performed by Novogene. Sequencing libraries were generated using the NEBNext® Ultra™ RNA Library Prep Kit for Illumina (NEB, USA). Clustering of index-coded samples was performed on a cBot Cluster Generation System using TruSeq PE Cluster Kit v3-cBot-HS (Illumina). Sequencing was carried out on an Illumina platform, and paired-end reads were generated. The filtered reads were aligned to the most recent version of the maize reference genome, B73_v4 (release-44), using HISAT2 (version 2.1.0, Kim et al., 2015). HTSeq v0.6.1 (Anders et al., 2015) was used to count the reads numbers mapped to each gene, and the Fragments Per Kilobase of transcript sequence per Million base pairs sequenced (FPKM) of each gene was calculated based on the length of the gene and reads count mapped to this gene. Differential expression analysis was performed using the DESeq R package (1.18.0, Anders and Huber, 2010), and p-values were adjusted using the Benjamini and Hochberg’s approach. Amino acid sequences of maize candidate genes were compared to Arabidopsis protein sequences using the BLAST tool (Altschul et al., 1997) on the TAIR website (https://www.arabidopsis.org).

### Statistical analysis

Statistically significant differences were determined by the statistical tests referred to in Figure legends using GraphPadPrism (version 4; GraphPad Software).

### Data availability

The raw RNAseq data will be deposited at NCBI SRT with SRA accession number xxx.

## List of author contributions

SM, IM, and LGS conceived the project and designed experiments. SM and MFV conducted most experiments. RB and IM performed GC-MS analysis of cuticle composition, and PS and SM performed cryo-microscopy. JC, NK, MAG and LGS helped to establish and conduct the leaf rolling assay. SM, RB, IM and LGS analyzed data. SM and LGS wrote the article with contributions of all authors.

## Funding information

This work was financially supported by U.S. National Science Foundation IOS1444507, a Deutsche Forschungsgemeinschaft (DFG) Fellowship (MA-7608/1-1) to SM, and by funding from the Canada Research Chairs program (CRC) to IM.

## Supplemental Data

**Supplemental Figure S1.** Bulliform strip number and width are not correlated.

**Supplemental Figure S2.** Positional analysis of epidermal cell shrinkage upon dehydration.

**Supplemental Figure S3:** Abaxial BC-like cells in bulliform-enriched mutants do not necessarily have cuticle nanoridges.

**Supplemental Figure S4:** Single compound wax profiles of bulliform-enriched cuticles.

**Supplemental Figure S5:** RNAseq analysis of bulliform-enriched mutants.

**Supplemental Figure S6:** Phylogeny of selected BAHD family acyltransferases with maize candidate genes.

**Supplemental Table S1.** Leaf rolling and bulliform strip patterning data

**Supplemental Table S2.** Genes differentially expressed in all three bulliform-enriched mutants

**Supplemental Table S3**. TAIR protein BLAST results for Zm00001d008957

**Supplemental Table S4.** TAIR protein BLAST results for Zm00001d050455

## Acknowledgements

We thank Prof. Anne Sylvester (University of Wyoming), Prof. Phil Becraft (Iowa State University), and Prof. Neelima Sinha (UC Davis) for donation of seeds. We also thank Prof. Siobhan Braybrook (UC Los Angeles) for use and help with the Keyence imaging system equipment. This work was financially supported by U.S. National Science Foundation IOS1444507, a Deutsche Forschungsgemeinschaft (DFG) Fellowship (MA-7608/1-1) to SM, and by funding from the Canada Research Chairs program (CRC) to IM.

